# Morphology as indicator of adaptive changes of model tissues in osmotically and chemically changing environments

**DOI:** 10.1101/2023.01.18.524301

**Authors:** Kevin Höllring, Damir Vurnek, Simone Gehrer, Diana Dudziak, Maxime Hubert, Ana-Sunčana Smith

**Author notes:** Corresponding author, ✉, (A. Smith).

## Abstract

We investigate the formation and maintenance of the homeostatic state in the case of 2D epithelial tissues following an induction of hyperosmotic conditions, using media enriched with 80 to 320 mOsm of mannitol, NaCl, and urea. We characterise the changes in the tissue immediately after the osmotic shock, and follow it until the new homeostatic state is formed. We characterise changes in cooperative motility and proliferation pressure in the tissue upon treatment with the help of a theoretical model based on the delayed Fisher-Kolmogorov formalism, where the delay in density evolution is induced by the the finite time of the cell division. Finally we explore the adaptation of the homeostatic tissue to highly elevated osmotic conditions by evaluating the morphology and topology of cells after 20 days in incubation. We find that hyperosmotic environments together with changes in the extracellular matrix induce different mechanical states in viable tissues, where only some remain functional. The perspective is a relation between tissue topology and function, which could be explored beyond the scope of this manuscript.

Experimental investigation of morphological effect of change of osmotic conditions on long-term tissue morphology and topology
Effect of osmotic changes on transient tissue growth behaviour
Analysis of recovery process of tissues post-osmotic-shock
Toxicity limits of osmolytes in mid- to long-term tissue evolution
Tissue adaptation to physiological changes in environment
Long-term tissue stabilisation under altered osmotic conditions.

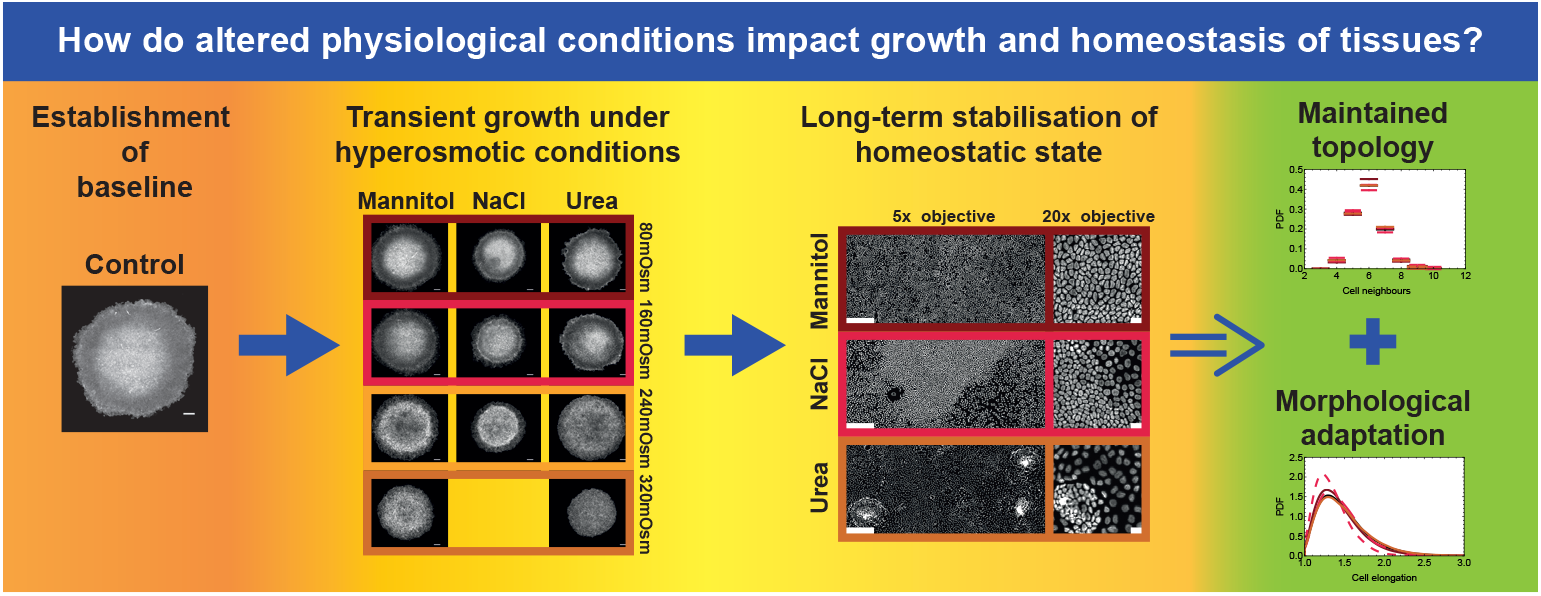

## 1. Introduction

Epithelial tissues can form, develop and survive in various chemical and mechanical environments while being exposed to different constraints and stresses [14]. These effects have been particularly well studied in the context of mechanoresponse, where changes in the tissues’ surroundings trigger an adaptation in terms of cell morphology [42, 29, 16, 15, 10], motility [28, 29, 43], proliferation rate [39, 12], and in the forces cells exert onto their neighbours as well as onto the extracellular matrix [10, 23]. Interestingly however, the structure of the tissue – reflected in its topology – was found to be preserved throughout changing conditions, as long as the tissue remains viable [15].

Besides adaptation to mechanical clues, tissues must respond to physiological changes in the osmotic background [46]. This is particularly important for renal cells, but the response mechanism was found to be preserved over a variety of other cells [38]. It was found that on short time scales hyperosmolarity involves cell shrinkage and changes in membrane tension, due to a rapid efflux of water across the plasma membrane [3, 35]. This typically induces an increase of the ionic concentration in the cytoplasm. To counter this effect on time scales of seconds to minutes, ion pumps in the plasma membrane, such as Na^+^-K^+^ - 2Cl^−^ co-transporter (NKCC), the Na^+^/H^+^-exchanger and the Cl^−^/HCO_3_^−^-exchanger start transferring small ions into the cell [20, 2]. Consequently, the osmotic pressure in the cytosol is raised and the cell starts to restore volume. However, the elevated ion concentration in the cytosol induces protein denaturation, DNA damages and molecular crowding [32], ultimately increasing the probability of cell death.

On the intermediate time scale of a few hours, the surviving cells start exchanging the small ions for amino acids using among others the amino acid transport System A [1], decreasing the ongoing damage to the cell. In the long-term response, complex signalling pathways are activated [38, 17], and amino acids are replaced by small organic osmolytes [46, 4], like betaine, myoinositol and taurine [44, 45]. If this stage is successfully reached, the cells have adapted to the changed conditions in terms of osmotic pressure and a new steady state should arise.

Most elements of the above adaptation process have been identified or confirmed to occur in several different cell types among which are the Madin-Darby Canine Kidney (MDCK) cells grown in various hyperosmotic conditions [34, 44, 45, 33]. Consequently, this cell line has been established as a good model system for studying general cell and tissue response to varied osmotic conditions. Therefore, MDCK cells were used to show that hyperosmolarity regulates endothelin release [37], and phospholipid biosynthesis [5].

Volume regulation in MDCK monolayers was found to directly couple to tension, where membrane reservoirs are used to compensate for swelling in hyperosmotic conditions. At the same time, a change in cell height was recorded upon different hyperosmotic treatments [30], strongly affecting the morphology of the cells’ apical membrane [41]. Changes in tension upon modification of osmotic conditions were also found to affect the selectivity in ion conductance [34], and paracellular transport through restructuring of tight junctions and localisation of claudin-2 and ZO-1 at cell-cell contacts [41]. Furthermore, since changes in tension are related to the tension of E-caderin bonds, osmotic conditions were found to correlate to lumen formation, cell proliferation, and the progression of the epithelial–to-mesenchymal transition [24]. This is then naturally reflected in a modulation of the cells’ migratory behaviour under osmotic stress [21].

Variations in osmolytes’ concentrations not only apply mechanical stresses on the cell through osmotic pressure [22, 6, 7, 9, 8], such variations also potentially interfere with processes controlling the structure of the tissue in homeostasis [11, 27]. However, the long term tissue adaptation emerging from the short-term response and the cooperative behaviour of cells has been poorly studied so far.

We here use MDCK-II colonies grown in various hyperosmotic conditions employing three different osmolytes: mannitol, NaCl, and urea. This allows us to compare the purely osmotic response induced by mannitol to effects that arise in physiological conditions (changes in ionic concentrations due to hydration levels and increased urea in the urinary system).

We first investigate the short-term dynamics of MDCK-II tissues grown in hyperosmotic conditions to determine concentrations of osmolytes which alter the tissues’ properties the most without being toxic. We then investigate the mid to long term dynamics of the tissue, where homeostasis is established in the entire colony. To understand the consequences of adaptation to modified osmotic conditions, we evaluate tissue topology and morphology, which have recently been shown to be very robust properties of a tissue’s homeostatic state [15].

We show that elevated osmotic conditions strongly affect tissue growth dynamics, and the properties of the homeostatic state. In the most concentrated viable osmolyte solutions, topological properties of the homeostatic state can be disrupted, especially with NaCl and mannitol, while urea has surprisingly little effects on tissue connectivity, instead leading to changes of transient behaviour and global tissue density.

## 2. Methods

### 2.1. Preparation of solutions used to create hyperosmotic conditions

Stock solutions of mannitol, NaCl and urea (all Roth, Germany) in DMEM/F12 (Gibco GlutaMax) with 5% FCS (Sigma-Aldrich) and 0.5% Penicilin/Streptomycin (Sigma-Aldrich) were prepared fresh every 4 days. Solutions were made in 50mL Falcon tubes by dissolving the necessary mass of respective powders initially in 35mL to 40mL of media and subsequently topping up with media up to the 50mL margin after the chemicals had successfully dissolved and the solutions had cleared.

### 2.2. Substrate coating and preparation

Non-sterile rectangular 18mm glass cover-slides were washed prior to coating with absolute ethanol, Milli-Q water and again ethanol. After drying under a sterile hood, the cover-slides were transferred to 6-well plates (Greiner BIO-ONE, Germany), and 500 μL of solution of 10 μgmL^−1^ collagen-I (Rat tail, Corning, USA) in 0.05M acetic acid (Riedel de Haën, Germany) was pipetted on top of the cover-slide without spilling to the rest of the well. Consequently, the lid of the 6-well plate was returned back and the cover-slides were left to coat overnight under the sterile hood. After (16±1)h of coating, the collagen solution was removed with a Pasteur pipette attached to a vacuum system until only a thin layer of liquid remained. A 300 μL droplet of PBS was then deposited on top of the cover-slide without spilling to the rest of the well. The procedure of PBS deposition and removal was performed once more followed by repetition of the procedure using DMEM/F12 (5% FCS) cell culture media leaving the cover-slides ready for cell seeding. As part of the well not covered by a cover-slide was left dry, the 250 μL droplet of media on top of the slide was sufficient for cell cultivation during cell attachment.

### 2.3. Cell preparation and seeding

We used the MDCK-II cell line (ECACC 00062107), obtained as a gift from Dr. Florian Rehfeldt (University of Bayreuth). The cells were cultured in a cell culture incubator that kept a humidified atmosphere of 95% H_2_O and 5% CO_2_ at 37 °C in T75 flasks in the presence of DMEM/F12 with 5% FCS medium. Seeding cells into the 6-well plates involved pipetting a 6 μL droplet of cell suspension containing 3 × 10^4^ cells, such that a clear separation between the deposited cells and the rest of the cover-slide was obtained. This ensured a uniform borderline of the cell monolayer once confluence was achieved about 12 h after seeding (no cells or cell islands were found outside the main cluster after confluence).

After 6 h to 8 h of incubation, the cell clusters were briefly inspected under the microscope. The wells with successfully developing colonies were washed once with DMEM/F-12 (5%FCS) medium which was concomitantly exchanged with 2 μL of fresh culture media containing either mannitol, NaCl or urea (80, 160, 240, 360 and 400 mOsm). For consistency, the medium exchange was also performed in control wells. Finally, the 6-well plates were transferred back to the incubator, where the medium was exchanged every 24 hours. Cells were fixed and stained for nuclei with Hoechst dye (Thermo Fisher, USA) and for F-actin with Phalloidin-TRITC (Sigma-Aldrich, USA) each day from day 1 to 4 (all conditions), as well as 20 days (only 240 mOsm) after seeding.

### 2.4. Image acquisition and analysis

The colonies were imaged on a Zeiss AXIO Observer.Z1 inverted microscope with a moving stage using a 5× objective, or a 20× objective. Images of entire colonies were stitched together from individual fields of view using ImageJ software [31].

To evaluate the density of cells in the colonies, cell nuclei were segmented, using the publicly available cellpose tool [40], with some custom modifications to reduce storage requirements for retaining the resulting segmentation. The results were visually validated, to confirm accuracy of the results. Some colonies were subjected to local thresholding to increase contrast and improve cellpose’s segmentation result. Under-segmentation errors were observed in regions of high density, where it was also hard to discern different nuclei visually. We estimate segmentation errors to be below 5%. Manual correction of this error does not change the results significantly.

From the pixels that were identified to belong to the same nuclei, the centre of mass (COM) position of the respective nuclei was then calculated. The centre of the colony was determined to be positioned at the arithmetic mean of the COM positions of all detected nuclei. Sorting the nuclei positions from lowest *d*_min_ to furthest distance *d*_max_ from the centre of the colony (based on our knowledge of the dimensions of a pixel), the range [0, *d*_max_] was then split into 30 intervals of equal size, each limited by a minimum radius *r*_low_ and a maximum radius *r*_high_. For each interval, the number or pixels *n*_pix_ visible within the field of view with a distance to the centre of the colony between *r*_low_ and *r*_high_ was then determined as well as the number of cell nuclei COM *n*_cell_ in the same range. Knowing the area *A*_pix_ of a single pixel, the mean cell density

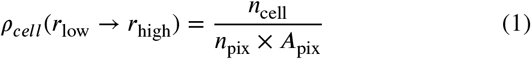

was then calculated and used as an estimate for the mean radial density profile for the central radius of the interval at 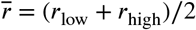

For the day 20 analysis, when the entire well is covered, the density analysis was repeated on a panorama of 5× images. However, instead of considering the distance of cells from the centre of the colony, the range [*x_min_*, *x_max_*] of x-coordinates of cell nuclei was instead dissected into 30 intervals, with the analysis for each interval again relying on counting the number of cells, the number of pixels and the denoted *x*-position of an interval being its centre *x* coordinate according to eq. (1).

### 2.5. Morphological and topological analysis of cells within the tissue

The morphological and topological analysis was performed on cells 20 days after seeding using 20× images. The cell shapes and connectivity was obtained through the Shape-based Voronoi Tessellation (SVT) algorithm already used in a previous publication [15]. We have shown that such a method is able to approximate the membrane area with an error up to 10%, the perimeter up to 5.5%, the elongation up to 10% and the neighbourhood up to 7%.

The statistical analysis of morphological information was performed on 3 independent clusters (4 in the case of NaCl) and the following number of cells have been recorded: 39 994 cells in control conditions, 67 180 cells in 240 mOsm mannitol conditions, 64 837 cells in 240 mOsm NaCl conditions, and 34 110 cells in 240 mOsm urea conditions.

Given the strong heterogeneity of density within NaCl-grown clusters, high and low density regions have been separated. Based on visual inspection of the images of one exemplary cluster, images with average density below 4200 cells mm^−2^ show low cell density everywhere and thus were used for the *low density* population. Contrary to that, images with average density higher than 7500 cells mm^−2^ were used for the *high density* population. The corresponding number of cells for the statistical analysis is 8724 cells and 14 352 cells respectively. This does not relate to the frequency of the high- and low-density regions occurring, as the high-density regions are generally rarer, but they do contain a far larger amount of cells in the same field of view.

Statistics for the cell area and perimeter are presented in dimensionless units. The cell area is divided by the mean cell area 〈A〉 within the respective cluster and the cell perimeter is divided by the square root of the same mean cell area 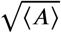. Therefore, cells from different clusters are divided by different mean values.

The Probability Density Functions (PDF) are built via the *SmoothHistogram* function provided by Mathematica. The function uses the default Gaussian kernel with a bandwidth which changes with the morphological measure considered. The bandwidth for the rescaled cell area is 0.05, for the rescaled cell perimeter 0.10, and for the cell elongation 0.05. Cell neighbourhood is based on a classical histogram with classes of size 1 centred on integer values.

## 3 Results and Discussion

### 3.1. Short-term response to hyperosmotic conditions

We first investigate the response to osmotic changes on a short time scale after cells have adapted to the new osmotic conditions at 24 h (see fig.1) and up to 48 h (see appendix fig.A9) after the initial shock, and still prior to the establishment of homeostasis.

**Figure 1:**
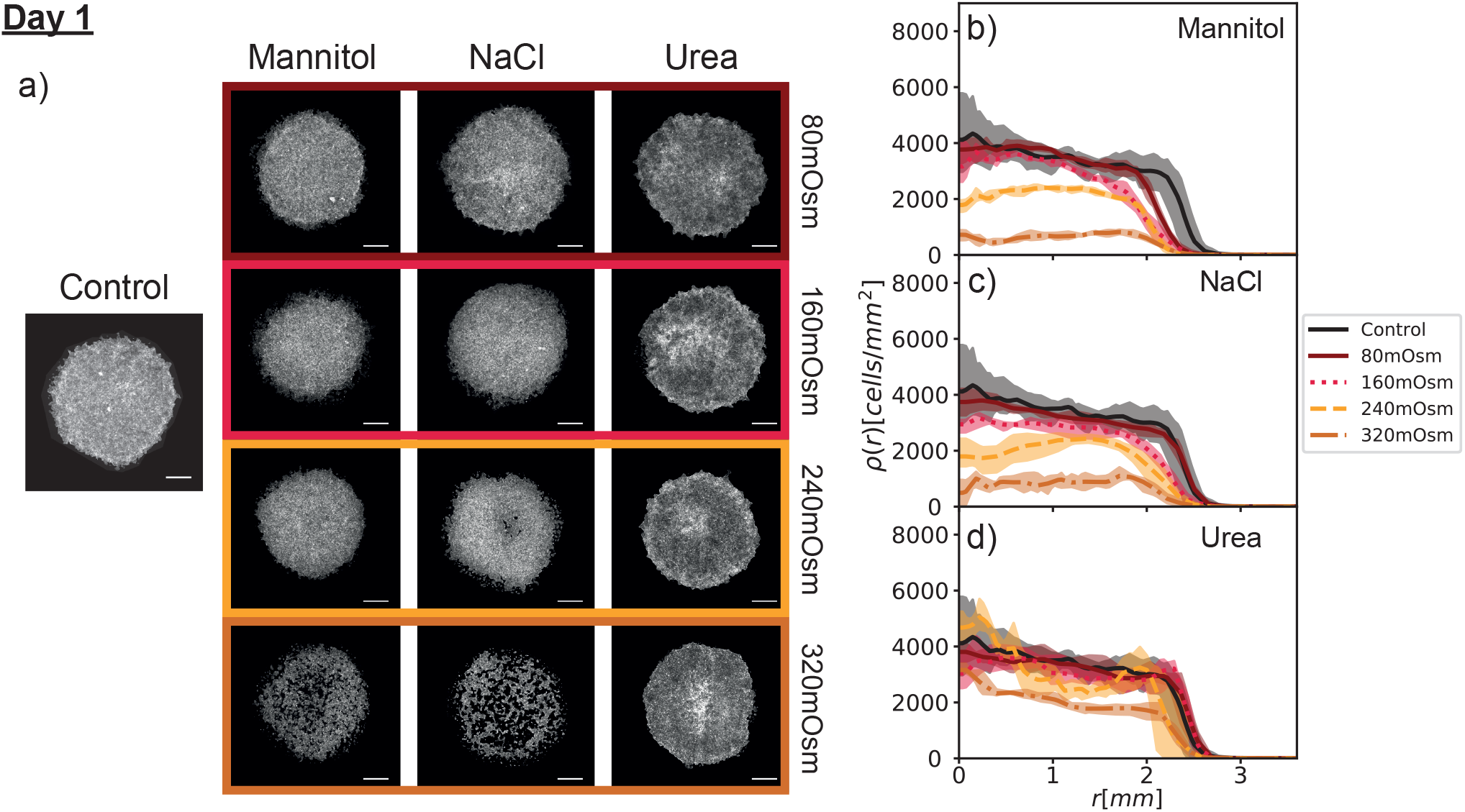
Actin images of day-one response to hyperosmotic conditions and associated nuclei density profiles. (a) 30 000 cells were seeded on collagen-I coated glass cover-slides and cultured for 24 h, fixed and stained. Light intensity in the images was enhanced such that all the clusters were visible (intensities are not comparable between clusters, yet faithfully depict qualitative differences within a single colony). Scale bars in the pictures represent 1mm. Density generally drops monotonously for increasing osmolyte concentrations. For 320 mOsm, increased cell death of the mannitol and NaCl colonies rendered them sub-confluent after 24 hours. Also, colonies at these concentrations are sensitive to cell handling and cells can detach completely if due care is not taken during media application. Nuclei density profiles for all osmolyte concentrations for (b) mannitol, (c) NaCl and (d) urea are shown. The black data in the respective graphs accounts for the control as a reference and are the same for all systems. Per osmolyte, *n* = 1 cluster was analyzed, split into four quadrants for error estimates. The plots includes the upper and lower bound of densities measured in the quadrants.

In these early stages, we find that our model tissues in 80, 160 and 240 mOsm are fully viable. At 400 mOsm conditions, all colonies detached within 15 min of the application of osmolyte enriched media and displayed crumbled cell debris. The concentration of 320 mOsm is in a crossover regime, with colonies treated with urea and mannitol being stable, while colonies subjected to NaCl detach on timescales of a few hours to a few days after the initial shock. This suggests that the cells under the NaCl stress are not capable of activating the intermediate response associated with the replacement of ions in the cytoplasm, and cell death becomes imminent. Furthermore, we overall observe the MDCK-II cells being less affected by urea than by mannitol and NaCl, likely due to their origin in the renal system.

We observe several generic responses of the model tissues. The most notable and systematic change is in the total cell number (see appendix table A1), which shows that the overall cell loss is increasing with the intensity of the osmotic shock. While at 80 mOsm only minimal changes can be observed for all three osmolytes (the biggest one being mannitol with about 20% loss), already at 160 mOsm of mannitol and NaCl, about 30% of cells did not survive the sudden change in the osmotic background. For 240 mOsm mannitol and NaCl, about 50% of cells died, an effect that is even more pronounced at 320 mOsm when the cell loss is about 80%. This is contrasted by urea where the cell death is less severe at about 20% and 45% at 240 mOsm and 320 mOsm, respectively. Importantly, the cells that do survive seem to restart their proliferation cycle, as the number of cells in all colonies double between day 1 and day 2 (table A1).

It is, furthermore, interesting to notice that cell death in immediate response to the change in hyperosmotic conditions did not appear homogeneously over the entire colony (see left panel fig.1). Evidence of this can be seen by inspecting the images of colonies taken within 18 hours after the shock. Specifically, we find for all concentrations of all osmolytes the edge retracts upon treatment albeit, for smaller osmolyte concentrations, the changes are within the statistical accuracy. Beyond 80 mOsm, the overall size of the colony is independent of the osmolyte concentration. This suggests that edge cells, many of which have leading cell morphology (i.e. larger size) and a higher rate of proliferation in the 0.5mm large compartment [15], are particularly sensitive to osmotic shocks, and detach more promptly.

Besides the very edge, cell death has more severe consequences in the centre of the colony, causing the colonies at higher concentrations to lose confluence there. Notably, the cells there seem to detach in patches. This is particularly acute for NaCl and mannitol. Given that the centre of the colony typically has a larger density prior to treatment, which amounts to smaller cell sizes and a lower rate of proliferation, this is consistent with the larger sensitivity of smaller cells (right panel fig.1). Notwithstanding, colonies subjected to urea at 240 mOsm and 320 mOsm concentrations do neither exhibit the same drop in cell density as other osmolytes nor the specifically lower centre density or a loss of confluence. Generally, central densities range from about 4000 cells mm^−2^, for control-colonies and colonies subjected to concentrations of 80 mOsm, down to 800 cells mm^−2^ at high 320 mOsm concentrations for mannitol and NaCl.

Already 48 hours after the osmotic shock, the number of cells in the colony nearly doubles (see appendix table A1), showing the recovery of the cell cycle. Consequently, the density profiles start to follow a normal growth pattern with the density increasing in the centre and the edge expanding. Average densities in the centre, however, shift up only slightly compared to day 1 with the highest densities at between 5000 cells mm^−2^ and 6000 cells mm^−2^ for some colonies subjected to low-concentration NaCl and the lowest being at 1000 cells mm^−2^ to 2000 cells mm^−2^ for high-concentration mannitol and NaCl systems. Again, the cells show a more pronounced resistance to urea as expected, but, starting from day 2, there is also a clear distinction between the densities observed under the influence of urea at and above 240 mOsm and the control.

### 3.2. Medium-term response and the establishment of the new homeostatic states

About 48 h to 72 h after the osmotic shock, we see a change in the general nature of density profiles throughout the colony, indicating a change in the growth behaviour of our model tissues. As expected, different compartments appear in the control: the central region where tissue is in homeostasis, the transition region with decreasing density, the actin enhanced stabilisation region, and the leading edge [15].

In hyperosmotic conditions of up to 160 mOsm, colonies also show a similar large-scale structuring but with much larger fluctuations across significant compartments which makes the profiles generally appear more jagged. This is likely an effect of the nature of a large number of cell divisions reacting to changes in the surroundings in a delayed manner, resulting locally in relatively high densities, which need some time to spread.

The fluctuations persist also at higher hyperosmotic conditions where the compartmentalisation is significantly affected, with the edge being poorly formed whereas the central compartment after 4 days reaches a homeostatic state (fig.2). Examples for this involve a transient dip in control density between the centre and the edge region and overshoots in colonies subjected to mannitol and NaCl at 80 mOsm. Similar effects are seen also in colonies treated with urea at 240 mOsm (see appendix fig.A10). This is likely an evolution of the slight excess initially observed on day 2 for those low concentration systems.

**Figure 2:**
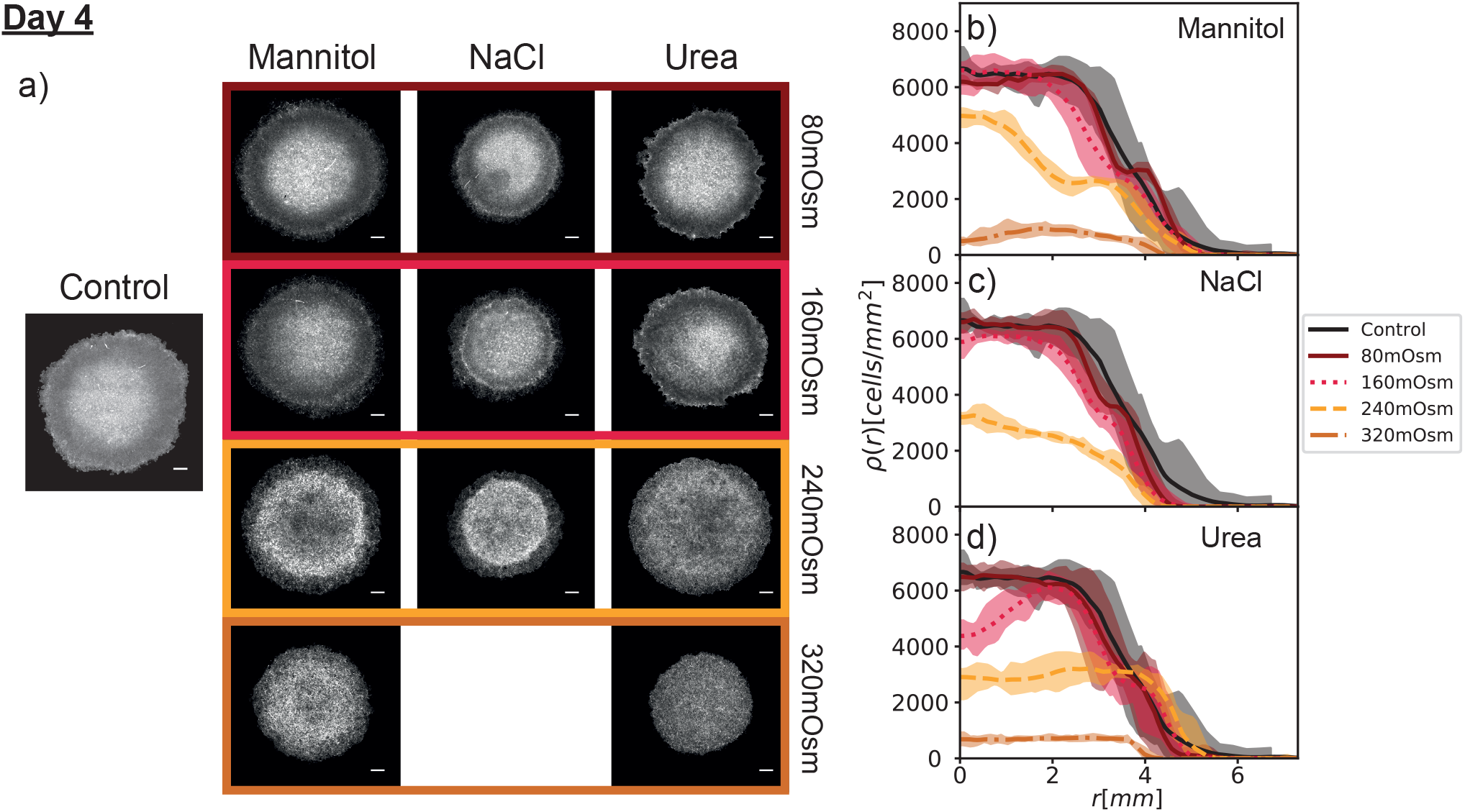
Actin images of day-four response to hyperosmotic conditions and associated nuclei density profiles. (a) shows the actin density profiles of representative colonies for the control and each combination of omsolytes and concentrations, obtained via the process described in fig.1. Here, *n = 2* cluster were evaluated for each combination, each split into quadrants for the radial analysis as described previously. Scale bars in the pictures represent 1mm. The graphs of the radial nuclei density profiles for (b)cmannitol, (c) NaCl, and (d) urea are shown in the same presentation as for day 1. All colonies of NaCl at 320 mOsm on day 4 had detached. Colonies at this concentration of NaCl at around this time were observed to be non-confluent, showed little to no clear polarisation and a generally turbulent structure. It was obvious that the colonies at NaCl-320 mOsm were not generally viable tissues on day 4.

While lower peak density and slightly reduced colony radius at higher concentrations persist up to the third day (see appendix fig.A10) and the fourth day (see fig.2), the colonies reach homeostasis. In the control and low concentration of all osmolytes, the homeostatic density is unaffected and takes values in the range of 6000 cells mm^−2^ to 7000 cellsmm ^2^ (see fig.2), consistently with earlier studies [15].

At higher concentrations (≥ 160 mOsm) the homeostatic state actually changes. This is evident through a strongly reduced density of the central region, which does not increase further in later days. Here, for the first time, we clearly observe differences between the three osmolytes. For example, urea-grown clusters at 240 mOsm achieve only 3000 cells/mm^2^.

Clusters at 320 mOsm settle at even lower densities. Mannitol - similar to NaCl - builds up density in the central compartment at a significantly reduced size compared to the control. However, it eventually settles into a low density state with a slightly expanded radius. Similar overshooting, but at smaller intensity is seen with urea and NaCl, the latter eventually inducing a complete tissue breakdown manifesting as detaching colonies and overall sub-confluent patches of cells remaining polarised to a low degree. This shows that persistent hyperosmotic conditions have a significant effect on the regulation of a tissue’s homeostatic state.

### 3.3. Resolving cell response to the osmotic shock from the cooperative tissue growth dynamics

#### 3.3.1. Modelling the establishment of homeostasis and steady expansion conditions

A number of the effects observed in the previously described experiments can be captured by a Delayed Fisher-Kolmogorov (DFK) formalism [13].Overall, tissue growth is presented in terms of two population densities *ρ_d_* (*t*) of dividing and *ρ_g_* (*t*) of growing cells forming the full cell density *ρ*(*t*) = *ρ_g_*(*t*) + *ρ_d_*(*t*), where *t* is time. The dependence on *t* will not be explicitly written out in the further discussion. The densities *ρ_d_*, *ρ_g_* and *ρ* evolve via a set of differential equations also depending on a time delay *τ* = *t* − *τ_d_*, where the delay is induced by the duration of the division phase *τ_d_* of the cell life cycle:

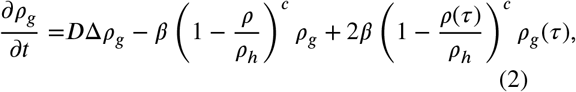

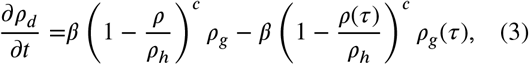

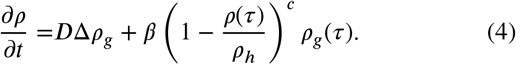

Furthermore, the target homeostatic density is *ρ_h_*, the growth coefficient is denoted by *β*, *c* is the mechanosensing exponent, and *D* is the diffusion coefficient modelling the tendency of cells to move into adjacent positions, i.e. active movement [19].

The effects of each parameters modified separately are shown in fig.3:

*ρ_h_*: This parameter represents the homeostatic density that the tissue will eventually converge to. It is mostly a scaling factor for the absolute target value of cell density and does not affect the qualitative radial evolution of the tissue.
*τ_d_* (first row in fig.3): This parameter is the time delay between a cell beginning division and it actually having split in two, amounting effectively for the mean duration of cell division. This introduces a delayed reaction of cell growth to the surrounding conditions and can lead to observations such as a density overshoot over the homeostatic density *ρ_h_* that is not spontaneously observed in Fisher-Kolmogorov equations without the time delay.
*c* (second row in fig.3): The mechanosensing exponent *c* in the logistic term (1 − *ρ/ρ_h_*)^*c*^ controls the shape of the relative growth term and its dependence on local cell density/pressure. It dampens or accelerates cell proliferation inhomogeneously as density approaches homeostasis. A higher value *c* ≫ 1 will lead to a softer shape of the tissue’s edge region as cell proliferation will slow down very quickly when approaching its target density *ρ_h_*, causing non-linear pressure profiles depending on density. The case *c* ≪ 1 on the other hand will lead to a constant ratio of dividing cells until very close to the homeostatic density, causing the tissue edge to be steep, thus well-defined. Such values of *c* also cause transient overshoots of density over *ρ_h_* which are usually located close to the tissue edge, in combination with the values for other parameters like *β* and *τ_d_*. In short, higher values of *c* lead to less overshooting and a smoother convergence to the target density, whereas lower values cause more overshooting and a steeper edge region. Both of these effects are related to faster and slower expansion-pressure density-scaling respectively.
*β* (third row in fig.3): The growth coefficient sets the time scale on which the tissue density converges towards its homeostatic density *ρ_h_*. It is directly associated with the proliferation pressure. Hence it is inversely linked to expansion pressure at the edge, with lower values of *β* causing faster expansion motion of edge cells and higher values triggering higher density buildup at the leading edge. As a result, it generally influences the width of the colony’s “edge area” together with the parameters *c*, *τ_d_* and *D*, and has an effect on tissue expansion. Additionally, it influences the tendency of tissue density to overshoot at the edge.
*D* (last row in fig.3): The diffusion coefficient in the radial direction is used to model cell locomotion triggered by the pressure gradient, but also active motion at the edge of the tissue. This global tissue property controls the speed at which the tissue expands and limits the extent to which purely local features can manifest.

**Figure 3:**
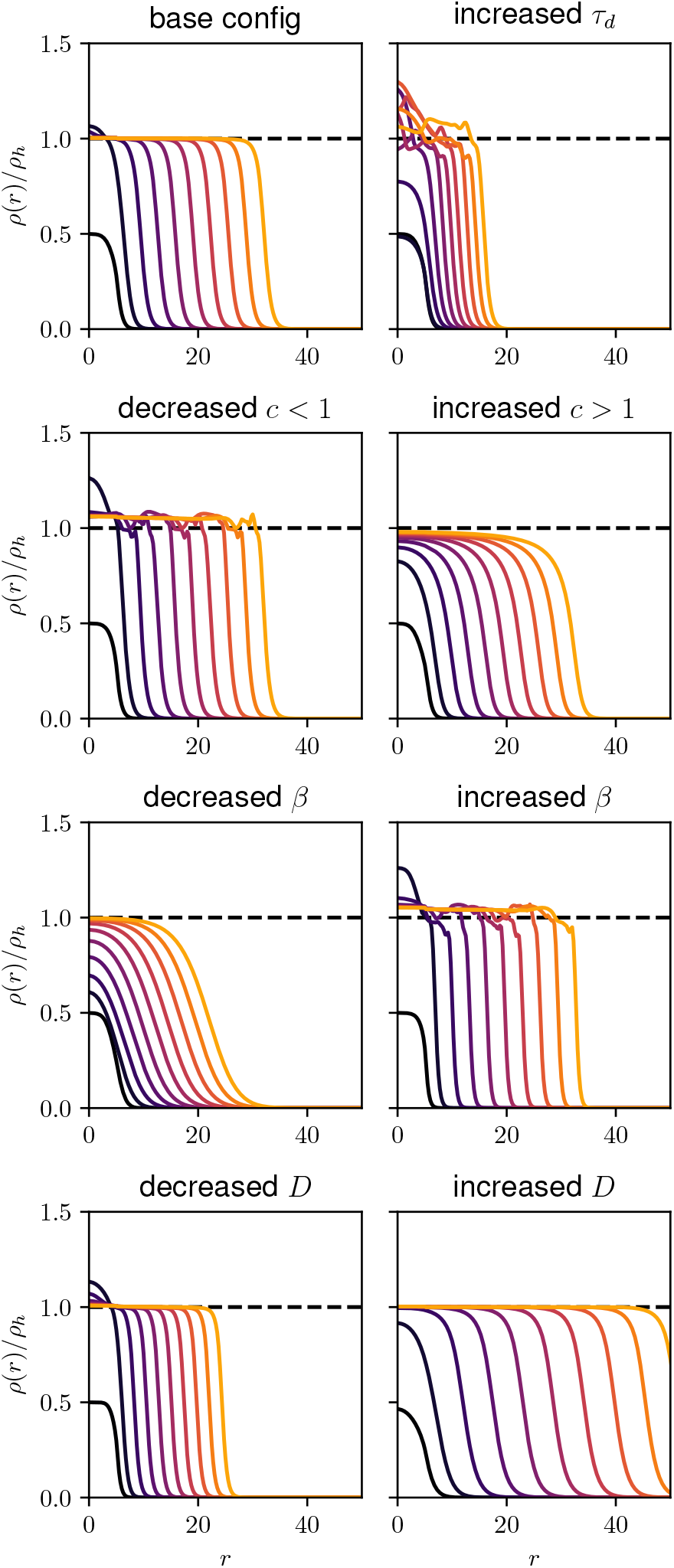
Characteristic effects of changes in parameters of the Delayed Fisher-Kolmogorov formalism on the time evolution for density profiles. As a base configuration, we picked *c* = 1, *D* = 1, *β* = 2, *τ_d_* = 1. The decreased values were *c* = 0.5, *D* = 0.5, and *β* = 0.3. The increased values were *c* = 2, *D* = 3, and *β* = 6 and *τ_d_* = 4. The colour gradient from dark to bright colours indicate the time evolution of the tissue density profile. Same colours indicate the same time during simulation across all plots. The dashed line is the configured homeostatic density *ρ_h_* = 1.0, which is included for visual reference. For context, we assess that the control’s parameters are somewhere in the environment of *ρ_h_* ≈ 6.5 × 10^3^ cellsmm^−2^, *τ_d_* ≈ 4.8 h, *c* ≈ 1.4, *β* ≈ 0.07 h^−1^, *D* ≈ 8.3 × 10^−3^ mm^2^h^−1^

To summarise, larger *D* and smaller *τ_d_* promote radial expansion of the tissue. Increasing *β* and *τ_d_*, or decreasing *c* affects the shape and width of the tissue edge region, and promotes transient oscillations and overshoots of the homeostatic density. They are tied to the speed of convergence towards the homeostatic density in competition with the parameter *D*, which spreads the width of the edge compartment.

#### 3.3.2. Evolution of the number of cells within the colony

In order to understand the dynamics of the tissue development, we first extracted the total number of cells in the colony as a function of time over 4 days of growth in different hyperosmotic conditions (see appendix table A1 for tabulated data). When such numbers are plotted relative to the number of cells on day 1 after the osmotic shock (fig.4) it becomes obvious that for all but systems subjected to 320 mOsm concentrations, the increase in the number of cells remains comparable to the control.

**Figure 4:**
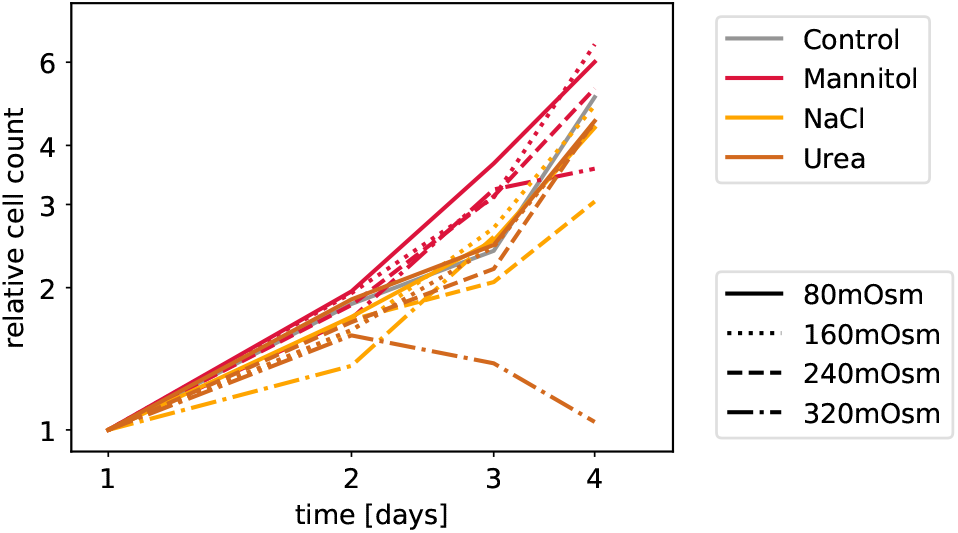
Evolution of the number of cells within the colony relative to day 1 cell numbers on a double-logarithmic scale. (see appendix table A1 for detailed data). We see a similar evolution for all involved systems, with only deviations, where the colony generally suffered from high concentration poisoning (i.e. NaCl and urea at 320 mOsm). All other combinations of osmolytes and concentrations did not deviate much from the control evolution, indicating overall similar proliferation cycle durations for all systems despite differing critical colony densities.

Small osmolyte-sensitive differences can, however, be noted. Namely, almost all colonies subjected to any concentration of mannitol exhibit larger relative cell counts compared to the control starting on day 3, which could indicate a change in proliferation parameters via a slightly decreased *τ_d_* or a reduction of the interval of in-between growth. On the other hand, both NaCl and urea at low concentrations remain consistently close to the growth curve of the control, indicating that the growth and division cycle has returned to normal parameters after the initial osmotic shock.

For concentrations of 320 mOsm for urea and NaCl, the cell count increases during the first 24 h just like in lower osmolyte concentrations. However, beyond that duration, the apoptosis takes over again, resulting in a drop of cell numbers, an effect that we cannot capture by the theoretical model that misses to explicitly account for apoptosis.

#### 3.3.3. Discussion of the measured growth profiles in the context of the DFK model

As discussed in previous works, the DFK model can capture the qualitative growth of the colony in standard conditions [13], which provides an excellent foundation for understanding the effects of an osmotically-modified environment on tissue growth dynamics. By comparison of trends observed in experiments, and in the DFK model, we see that foremost, an increase in osmolyte condition causes a reduction in the homeostatic target density *ρ_h_* across the board with the precise density vs hypertonicity dependence being tied to the type of osmolyte being applied.

From the cell count statistics across colonies submitted up to 240 mOsm, we infer that the division time *τ_d_* does not significantly change after recovery from the initial shock, except perhaps slightly in mannitol. The mechanosensitivity parameter *c* does not generally seem to change a lot in lower concentrations (≤ 160 mOsm), only at 240 mOsm small changes are arguably present. This indicates that from the standard environment until harsh, almost toxic conditions, tissues preserve the relation between the effective tension experienced by the cell and the cell proliferation process.

The convergence parameter *β*, however, seems to increase when colonies are subjected already to low concentrations of osmolytes. This is possibly a consequence of the recovery process aiming to restore the retracted leading edge, which requires an increase in proliferation pressure. A good example are colonies subjected to NaCl at 240 mOsm. These colonies initially overshoot its target density *ρ_h_*, due to a delay in proliferation, but then tissue expansion is triggered to smooth out this overshoot and spread the tissue to a more appropriate size. Since there is no clear leading-edge bump appearing during the overshooting, this points to an increase in *β* compared to the control, and not a change in *c*.

Changes to *β* are to different degrees accompanied by variations in motility *D* depending on the hyper-osmotic conditions. Towards higher concentrations of osmolytes, *D* seems to increase. This may be, in part, promoted by overall lower concentrations of cells in the tissue, which remain in a highly active state. The result is however, the smoothing out of features that readily appear at lower osmolyte concentrations. For example, at 240 mOsm of a mannitol-treated tissue, the edge is much wider and less steep, hinting at a reduced growth parameter *β* and an increase in *D*. The convergence to the target density is still very quick, which may then be rooted in the fact that the tissue initially was already very close to this new target density. Still, keeping a similar convergence speed towards the target density, an increase in *D* would need to be compensated by an increase in *β*, which in combination explains the faster edge expansion while converging towards the target density faster and without significant overshoots.

### 3.4. Long-term adaptation and structural changes within the tissue

We finally study the long term adaptation of tissues grown in environments with elevated osmolyte concentrations. To observe the most significant differences between systems, we restrict our investigation to concentrations of 240 mOsm.

We focus on characterising the properties of the homeostatic state, as the tissue is now covering the entire well. We first analyse the cell density and fluctuations after 20 days of incubation (see appendix table A2 for precise values and ranges). At this point, not only the adaptation to hyperosmotic conditions should have taken place, but also the mechanical environment should have been adjusted, as the cells have replaced the initial collagen matrix with the one secreted by themselves. The aim is to elucidate the joint effect of mechanical and osmotic adaptation, in an osmolyte-specific fashion.

**Figure 5:**
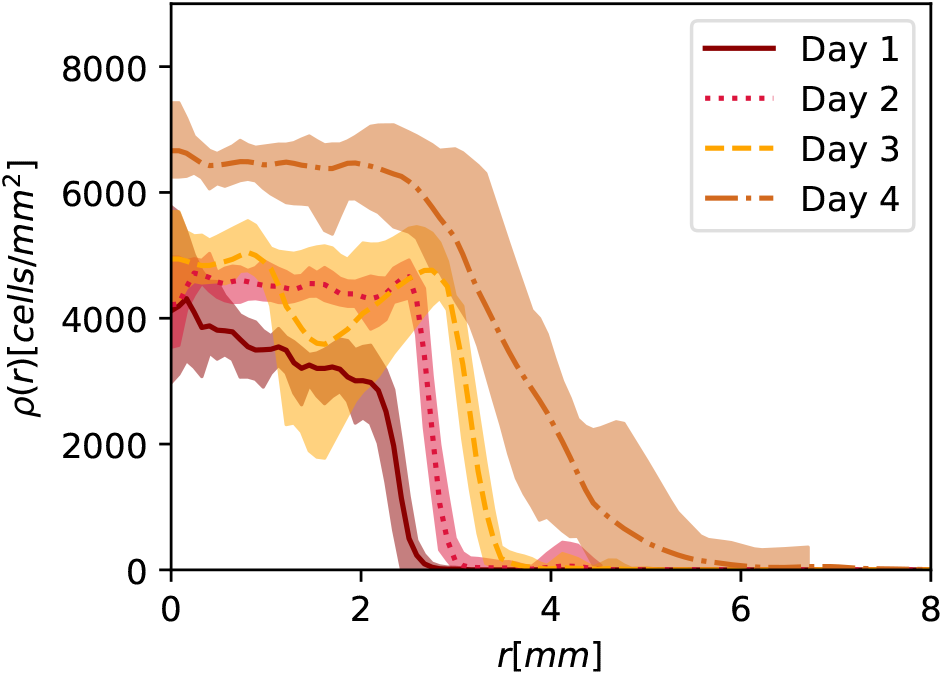
Time evolution of the density profiles from day 1 to day 4 for the control colonies.

#### 3.4.1. Changes in the macroscopic properties of the homeostatic state

For the control, long-term mechanical adaptation results in a decrease of the homeostatic density, as described previously [15], from about 6500 cells mm^−2^ measured on day 4 to (4669 ± 352) cells mm^−2^. Similar effects are seen in tissues grown with either urea or mannitol, however, the two conditions produce different homeostatic densities (see fig.7b-e). Urea for one, forces low density states of (3834 ± 422) cells mm^−2^, whereas mannitol pressurises the tissue towards high density homeostatic states of (5290 ± 470) cells mm^−2^. Interestingly, strong density fluctuations appear in a typical control sample (see top panel fig.7a) with densities reaching values between 3210 cells/mm^2^ and 5875 cells/mm^2^. Notably, the high-density regions in control samples are rather small and spread far apart. Fluctuations of a similar magnitude but with smaller characteristic length scales could also be observed in the mannitol-treated tissues. To the contrary, in urea the fluctuations are nearly suppressed, with only small islands of very high density appearing within the domes (see fig.7a). Notably, these domes are present in the control but much smaller, whereas they are nearly non-existent in tissues subject to mannitol or NaCl.

**Figure 6:**
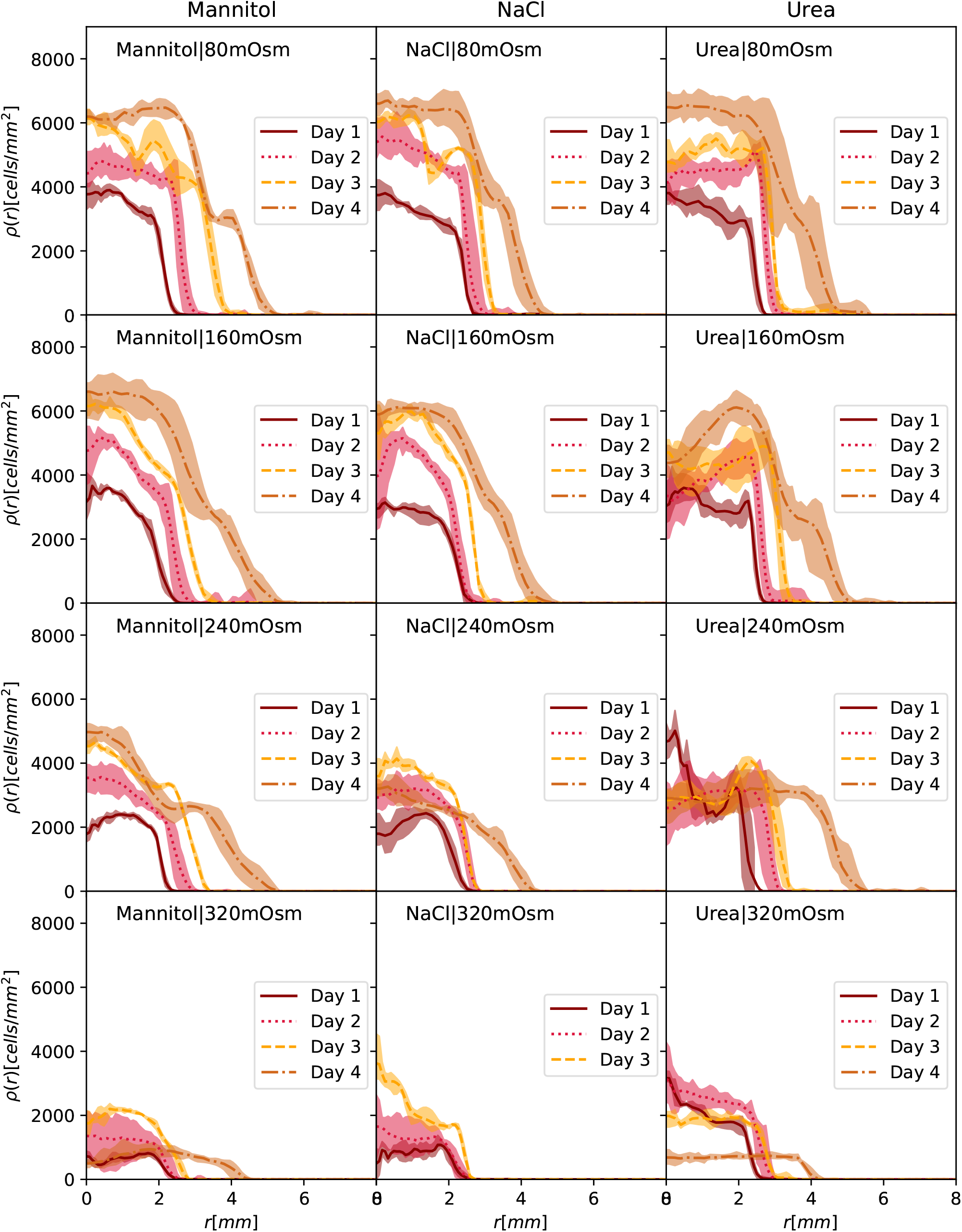
Time evolution of the density profiles from day 1 to day 4 for colonies submitted to different osmolytes (mannitol, NaCl, urea) at different concentrations. (80 mOsm, 160 mOsm, 240 mOsm and 320 mOsm).

**Figure 7:**
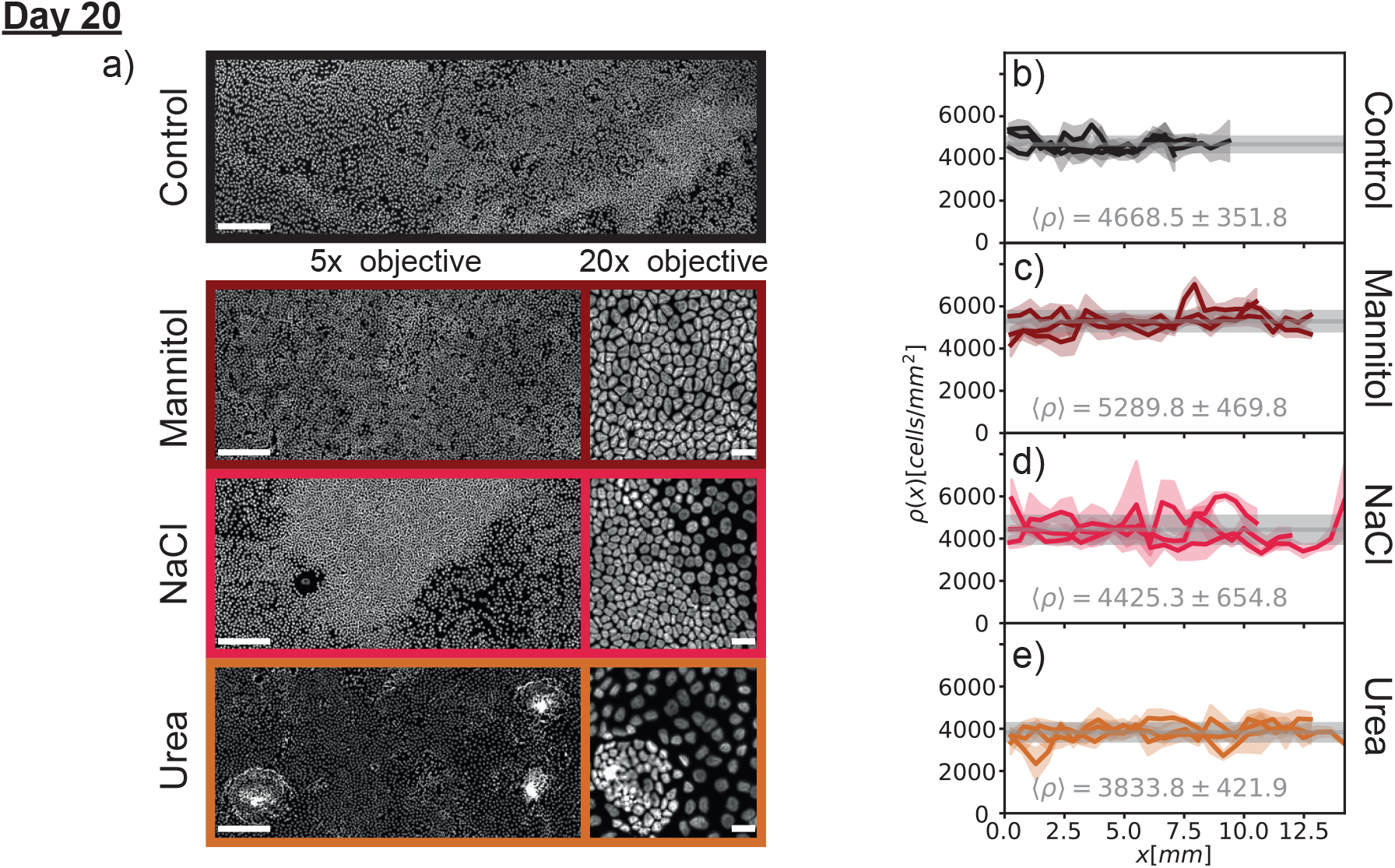
Visual characteristics and density profiles of steady state MDCK-II colonies at 240 mOsm on day 20. Epifluorescence images of the MDCK-II tissues grown in solution characterized with different osmolytes in hyperosmotic conditions: (a) Control conditions, (b) uniform density in mannitol, (c) heterogeneous density domain formation in the case of NaCl, and (d) frequent dome formation in the case of urea. Cell culture, staining and imaging were performed as described in the method section. The left side shows both 5x pictures (scale bar right 250 μm) and 20x pictures (only for osmolitic conditions, scale bar left 25 μm) whereas the right hand side of the figure shows characteristic density profiles for one representative colony for each osmolyte and control. For the density profiles, the cell nuclei were again segmented in stitched panorama 5x images and grouped and discretized by their *x*-coordinate instead of their radial position relative to the center of the colony. The gray horizontal line and the surrounding gray area visualize the mean density and the range of standard deviation of all colonies.

For NaCl, the average homeostatic state remains very similar to that of the control ((4425 ± 655)cells/mm^2^ and (4669 ± 352)cells/mm^2^ respectively). However, the length scale of density fluctuations becomes so large that, while still rare events, macroscopic regions of very high density occur in the NaCl-subjected colonies surrounded by a clearly distinct low-density state (see fig.7a for a high-density region in NaCl).

#### 3.4.2. Morphological and topological changes in the homeostatic state

It was previously shown that the topology of epithelial tissues is a property that is highly conserved, as characterised by the distribution of the number of cell neighbours [15, 36]. Furthermore, it was shown that changes in the mechanical properties of the extracellular matrix affect the density, but not the normalised morphology of cells comprising the tissue [15]. Therefore, a characteristic, stable and representative measure of the tissue state in terms of normalised area, and perimeter distributions could be found for the samples equivalent to our control. We here ask the question of the conservation of these measures in the scenario of strong alteration of osmotic conditions.

While typically cell area distributions are mono-modal including the one of the control, NaCl clusters show two states and accordingly a bimodal statistic in fig.8a. We therefore isolate and characterise each state independently (see *methods* 2.5). For the rescaled area and rescaled perimeter distributions (fig.8b and c), we find that for most populations of cells, the distributions fall onto the same curve. The exception are the distributions associated with tissues grown in mannitol as well as the high density state developing with NaCl. Both their distributions are narrower which hints towards a more homogeneous tissue, a conclusion which is corroborated by visual inspection of the cluster, as shown in fig.7. These exceptions are not translated to the distributions of cell elongation (fig.8d), where the clear outlier is the distribution of the low density state of the NaCl-grown cluster. These cells appear to be more circular on average.

**Figure 8:**
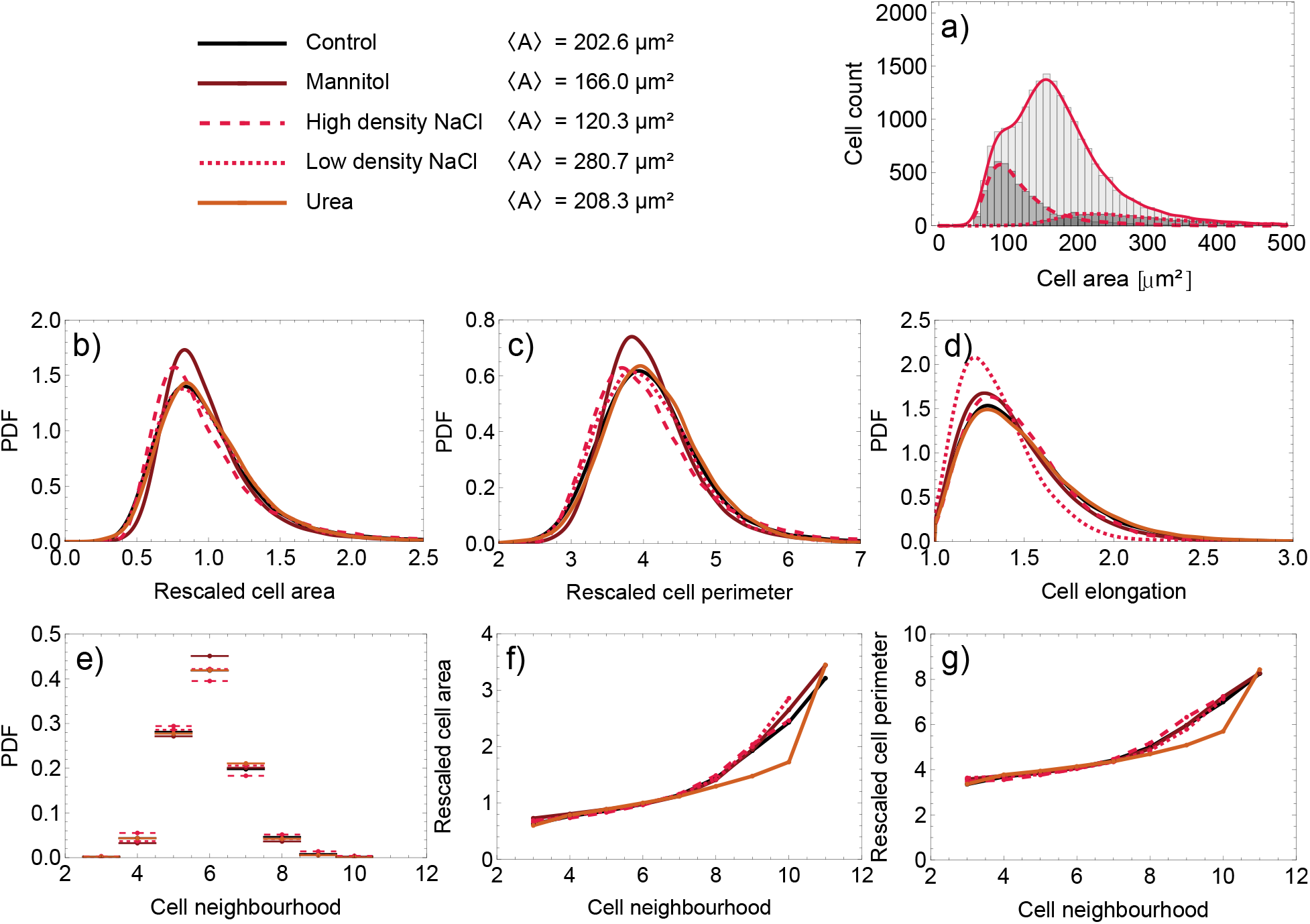
Self-similarity of morphological and topological distributions for different osmotic conditions. (a) Illustration of the separation of the global NaCl into its two sub-populations at high and low density by considering an histogram of the cell count. Only one cluster is considered for the illustration. (b,c,d,e,f,g) Probability Density Function (PDF) of the rescaled cell area (b), rescaled cell perimeter (c), cell elongation (d), cell neighbourhood (e), Lewis’ law (f) and Desh’s Law (g). All histograms have been prepared as described in section 2.5.

Further insight into the morphology of the tissue is given by the Lewis and Desh laws (fig.8f and g respectively), which relate the mean area and perimeter of cells respectively to the amount of neighbours those cells have. For example, in statistics of tessellations with Poisson-distributed generators, the perimeter and the area linearly increase with the number of neighbours - a larger cell has a proportionally larger number of proximate cells [18]. For tissues, this was found not to always be correct, as for example in our control samples [15]. Here, we find for all four conditions, as well as when separating the two populations obtained for NaCl-grown clusters, basically a universal, albeit nonlinear dependence. Some deviations can be observed for urea for 9- and 10-sided cells, but the limited statistics (184 and 16 cells respectively out of 34110) do not allow for any significant conclusion of alternate structuring behaviour.

We finally study the topology of our tissue, characterised by the distribution of the number of neighbours (fig.8e). These distributions by and large do not seem to be affected by the application of the osmolytes. There is no coupling to variations in the distributions of elongations. However, tissues developed in mannitol-rich medium and the high cell density state under NaCl show different characteristics. This is consistent with the deviations in the normalised area- and perimeter-distributions.

Tissues under mannitol are significantly more hexagonal, with an increased number of 6-neighbour cells. The high density state under NaCl is much less hexagonal compared to mannitol but also the control, displaying a significant surplus of cells with 5 and 4 neighbours and a depletion of cells with 7 neighbours. This is the first indication that it is actually possible to alter the topology of the tissues. The consequence of the change remains to be understood. Yet it is interesting to notice that neither of these samples present domes, which are an indication of trans-epithelial transport, obviously impeded in these tissues.

All other tissues, which are able to restore the morphological and topological properties equivalent to that of the control, form domes as a consequence of their regular tissue activity. The domes may be very different in frequency and sizes but show that these tissues are still functional. It is perhaps not so surprising that tissue subjected to urea does fall into this class, likely because of the origin of MDCK-II cells in the renal systems.

### 4. Conclusions

In this article we focused on characterising the fast and intermediate response of model tissues to a hyperosmotic shock induced by different concentrations of mannitol, NaCl and urea. We then observed the adaptation of the tissue to permanent but non-toxic hypertonic environments. We report on several important findings:

- As an “instantaneous” consequence of the osmotic chock, we find that the cell-survival within the model tissue depends on the intensity of the osmotic shock but also on the local environment. Particularly sensitive seem to be leading edge cells, and cells in the interior of the cluster, which represent either extremes of the size-, density- and proliferation rate spectra. We therefore suspect a connection to the deviation of cell sizes from an ideal configuration or a link to the pre-shock proliferation profiles. Either of these theses require more thorough investigation and should be the focus of future research.
- Independently of the intensity of the hyperosmotic shock, the cell cycle is restored relatively quickly within a day, with the numbers of cells nearly doubling after 48 hours. Furthermore, from the analysis of the growth patterns, we deduce that the relation between cell proliferation and effective tension in the tissue is unaffected until nearly toxic conditions.
- Despite being able to restart the cell cycle and nearly double the cell count daily in the short-term, the tissues treated with large concentrations of NaCl die on intermediate timescales of 3-4 days due to inabilities to adjust to persistently strong ionic conditions.
- On the intermediate time scales, we find evidence that both the cooperative motility of the cells and the pressure induced by proliferation are affected by highly hypertonic environments, compared to the control.
- The strong density fluctuations in growing tissues point towards the fact that fundamental features of MDCK-II cells’ tissue development are not altered on the level of single cells, but on the level of coordinated cell behaviour as a consequence of changes in boundary conditions, and the time-delays between density adjustment and the onsets of cell division.
- A hyperosmotic environment clearly decreases the transient density of the homeostatic state in the short to intermediate time frame and affects density fluctuations in an osmolyte-sensitive manner.
- On long time scales, adaptation of the tissue to high osmolyte concentrations, which also involves changes in the mechanical environment, further reduced the density of the homeostatic state. Interestingly, the effect of this adaptation is smaller in mannitol-treated tissue than in the control, such that its density after 20 days in a hypertonic environment is higher than the control. In the control, the MDCK-II cells exhibit strong fluctuations between regions resulting in high- and low-density domains, the former however, being very local. For NaCl, the bi-phasic organisation is stabilised and two different homeostatic states are found to coexists. Importantly, not only that the two states differ by average density, but they are also different to one another from cells’ topological and morphological viewpoints. This suggests that these two phases are really different mechanical states of the tissue.
- Like in NaCl, a modified homeostatic state was found for tissues subjected to high concentrations of mannitol. Urea, on the other hand retains the same topology and morphology as the control, suggesting that the mechanical state is preserved. To contrast others, these tissues remain functional, executing a strong apical to basal trans-epithelium transport, which opens an interesting question about the relationship between preservation of the mechanical state and tissue functionality.

To conclude, we find that tissues as a whole display different patterns of response to hyperosmotic conditions, which in turn affect the cooperative cell motility and proliferation pressure. Even if the cell cycle is conserved, and development of a tissue is not impeded, the homeostatic state is strongly affected, particularly in harsh conditions induced by high concentration of salt and mannitol. The results are viable tissues with reduced functionality, which is, interestingly, accompanied by changes in the mechanics of the homeostatic state.

The findings of our paper could have important implications in a physiological context. It has been, namely, established that a high salt diet is associated with accumulation of Na ions in the skin, over long periods of time [25]. This induces an adaptation processes in the skin to hyperosmotic conditions, which also affect skin function and wound healing [26]. Our data would suggest that the effects of hyperosmotic conditions could be reflected in the structure of the tissue - the morphology of the cells and their connectivity. These parameters could be explored as markers of a condition prior to the actual onset of a patological response. Further investigation is, however, necessary to uncover the relation between the hypertonicity and the mechanical state of homeostatic tissue on the level of molecular biology, a task that we hope will be addressed in future.

## CRediT authorship contribution statement

**Kevin Höllring:** Conceptualization, Methodology - Developed the Model, Software, Formal analysis, Writing - Original Draft, Visualization, Data Curation. **Damir Vurnek:** Investigation, Execution of experiments, Validation, Writing - Original Draft (methods), Visualization. **Simone Gehrer:** Formal analysis. **Diana Dudziak:** Resources, Lab oversight. **Maxime Hubert:** Formal analysis, Writing - Original Draft, Visualization, Data Curation. **Ana-Sunčana Smith:** Conceptualization, Supervision, Data Curation, Writing - Review & Editing, Project coordination.

## 5. Acknowledgements

This work was in part financially supported by the Deutsche Forschungsgemeinschaft (DFG) through the collaborative research center SFB TRR 305 - B05. We, furthermore, acknowledge funding by the DFG “Mechanobiology in Epithelial 3D Tissue Constructs” - 363055819/GRK2415 and the intramural funds by the IZKF, project A80. Finally, we benefited from contributions of the Emerging Fields Initiative “BigThera”, which was partly supported by the Staedler Foundation. We thank the Optical Imaging Center Erlangen (OICE) for their support.

## A. Appendix

**Figure A9:**
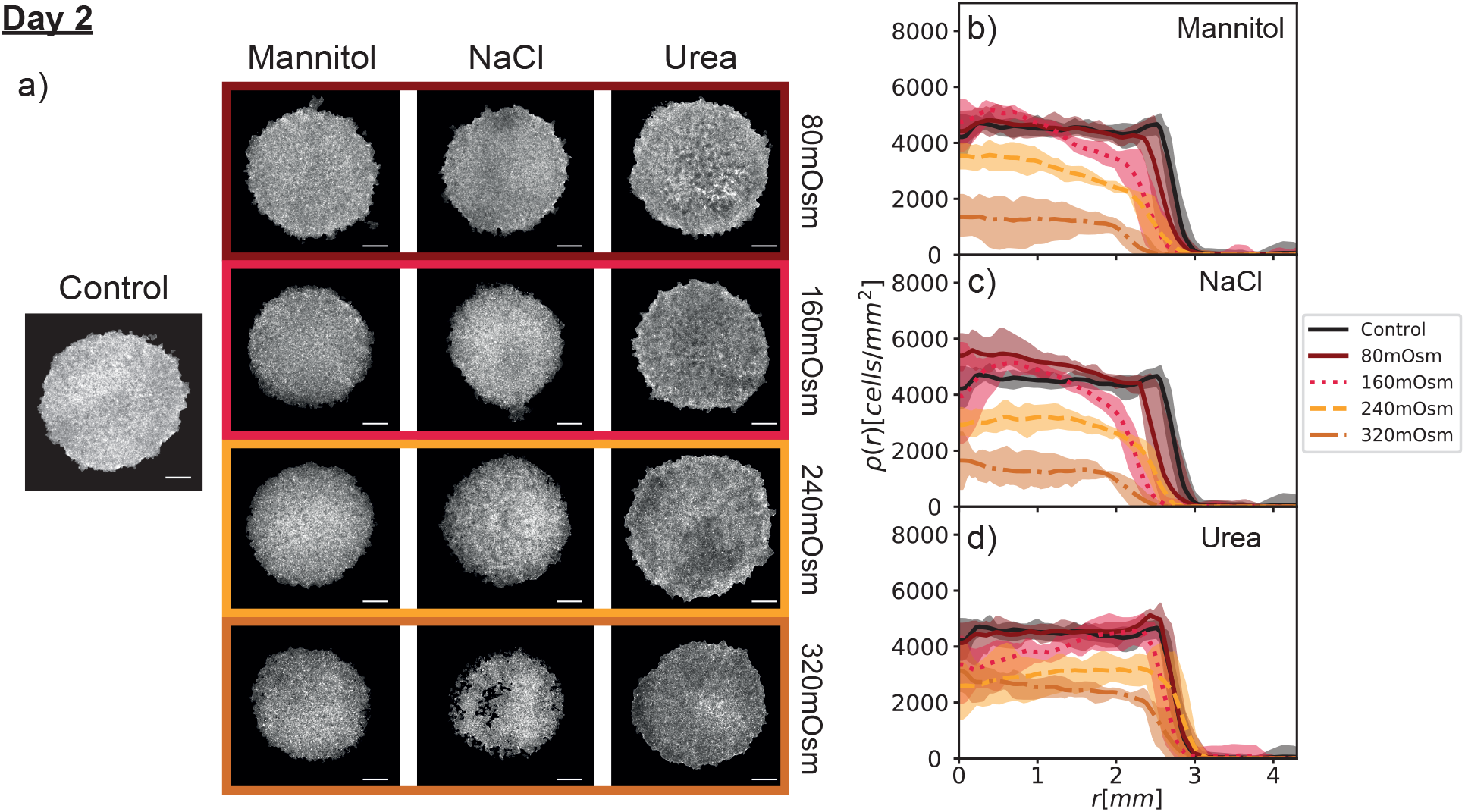
Actin images of day-two response to hyperosmotic conditions and associated nuclei density profiles. (a) shows the actin density profiles of representative colonies for the control and each combination of omsolytes and concentrations, obtained via the process described in fig.1. Scale bars in the pictures represent 1mm. Here, *n* = 2 clusters were evaluated for each combination, each split into quadrants for the radial analysis as described previously. The graphs of the radial nuclei density profiles for (b) mannitol, (c) NaCl, and (d) Urea are shown in the same presentation as for day 1

**Figure A10:**
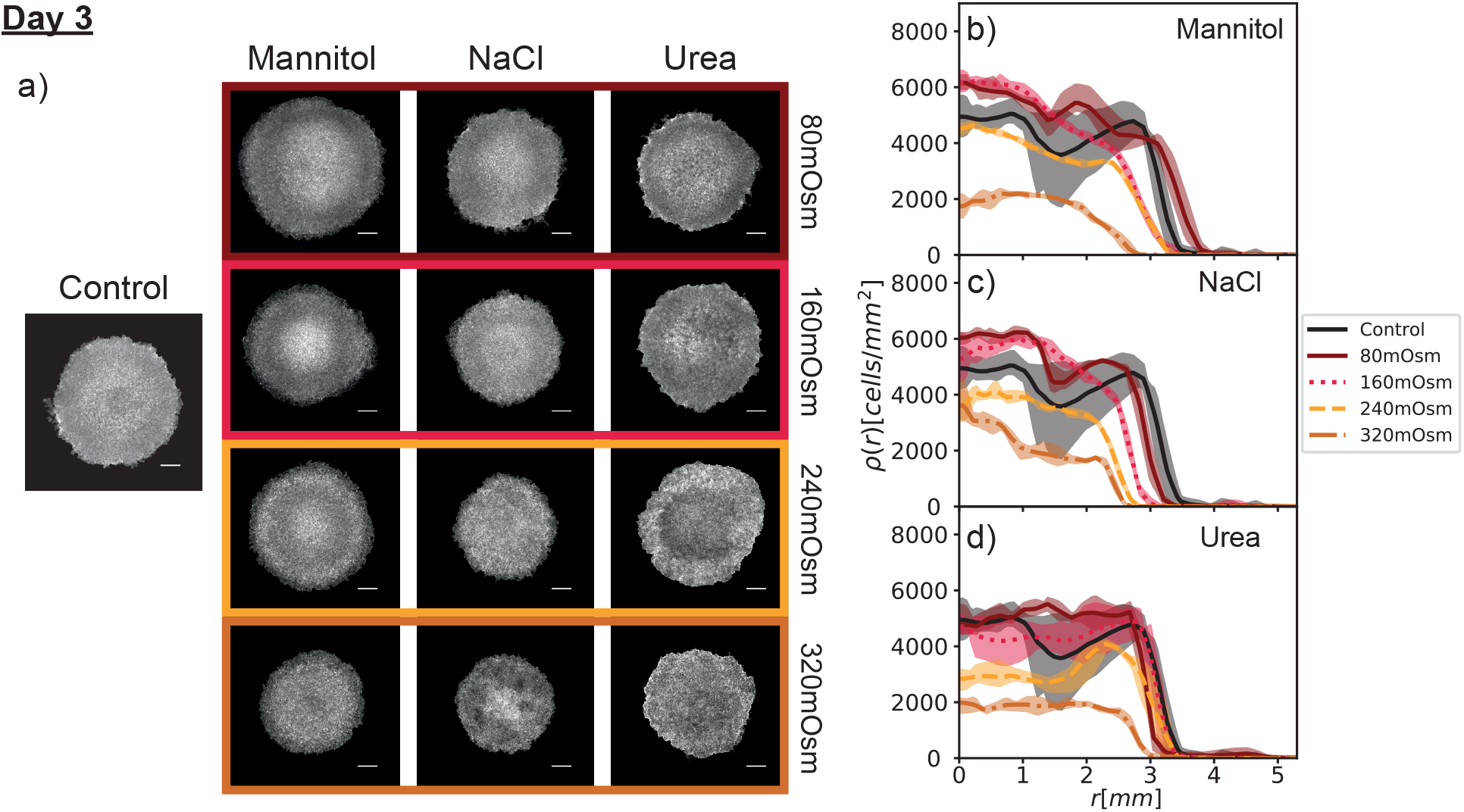
Actin images of day-three response to hyperosmotic conditions and associated nuclei density profiles. (a) shows the actin density profiles of representative colonies for the control and each combination of omsolytes and concentrations, obtained via the process described in fig.1. Scale bars in the pictures represent 1mm. Here, *n* =1 cluster was evaluated for each combination, each split into quadrants for the radial analysis as described previously. The graphs of the radial nuclei density profiles for (b) mannitol, (c) NaCl, and (d) urea are shown in the same presentation as for day 1

**Table A1.** Mean number of cells per colony on days 1 to 4. The number statistics of segmented cells per colony on days 1, 2, 3 and 4, 24 h to 96 h after the osmotic shock obtained via integration of the radial density profiles, rounded to the nearest integer.

| Day 1 |  |  |  |  |
| --- | --- | --- | --- | --- |
| Control |  | 58656 |  |  |
| Concentration |  | Mannitol | NaCl | Urea |
| 80 mOsm |  | 48324 | 56043 | 59007 |
| 160 mOsm |  | 37859 | 42108 | 57488 |
| 240 mOsm |  | 29330 | 32172 | 46167 |
| 320 mOsm |  | 10571 | 13882 | 32132 |
| Day 2 |  |  |  |  |
| Control |  | 108351 |  |  |
| Concentration |  | Mannitol | NaCl | Urea |
| 80 mOsm |  | 94920 | 97215 | 111452 |
| 160 mOsm |  | 73713 | 69171 | 93564 |
| 240 mOsm |  | 53901 | 55049 | 78049 |
| 320 mOsm |  | 18186 | 18970 | 50923 |
| Day 3 |  |  |  |  |
| Control |  | 140384 |  |  |
| Concentration |  | Mannitol | NaCl | Urea |
| 80 mOsm |  | 176990 | 141108 | 145304 |
| 160 mOsm |  | 117511 | 112653 | 140312 |
| 240 mOsm |  | 91579 | 66121 | 101171 |
| 320 mOsm |  | 34117 | 35572 | 44446 |
| Day 4 |  |  |  |  |
| Control |  | 296772 |  |  |
| Concentration |  | Mannitol | NaCl | Urea |
| 80 mOsm |  | 290317 | 244309 | 265761 |
| 160 mOsm |  | 247709 | 204469 | 256342 |
| 240 mOsm |  | 154824 | 97882 | 208050 |
| 320 mOsm |  | 37750 | - | 33409 |

**Table A2.**
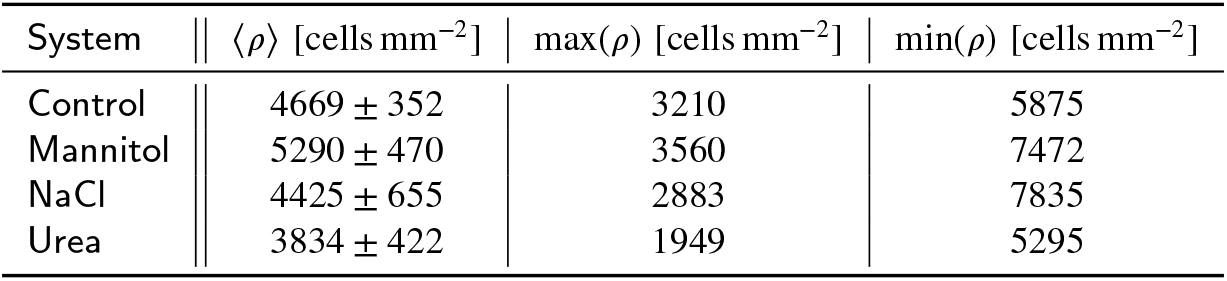
Day 20 saturation density. Mean density on day 20 of the 240 mOsm concentration for mannitol, Urea and NaCl as well as the Control system with standard deviation error estimate after the full well has been covered by the growing colony. Also presented are the local minimum density and the local maximum density. Density was calculated by slicing the panorama x5-microscopy image into slices along the z-direction after detecting nuclei positions via cell-pose in a local-threshold image produced by ImageJ after stitching with the appropriate plugin. The density was then calculated from the number of cell nuclei visible in each slice divided by the number of visible pixels, using the dimension *l* = 1.253 μm of the square-shaped pixels.

| System | \|\| $\langle \rho \rangle$ [cells $\text{mm}^{-2}$] | max($\rho$) [cells $\text{mm}^{-2}$] | min($\rho$) [cells $\text{mm}^{-2}$] |
| --- | --- | --- | --- |
| Control | \|\| 4669 ± 352 | 3210 | 5875 |
| Mannitol | \|\| 5290 ± 470 | 3560 | 7472 |
| NaCl | \|\| 4425 ± 655 | 2883 | 7835 |
| Urea | \|\| 3834 ± 422 | 1949 | 5295 |

## Notes

### Competing Interest Statement

The authors have declared no competing interest.

## References

[1] Alfieri, R.R., Bonelli, M.A., Petronini, P.G., Desenzani, S., Cavazzoni, A., Borghetti, A.F., Wheeler, K.P., 2005. Hypertonic Stress and Amino Acid Deprivation Both Increase Expression of mRNA for Amino Acid Transport System A. Journal of General Physiology 125, 37–39. URL:https://rupress.org/jgp/article/125/1/37/44192/Hypertonic-Stress-and-Amino-Acid-Deprivation-Both, doi:10.1085/jgp.200409195.

[2] Alfieri, R.R., Petronini, P.G., 2007. Hyperosmotic stress response: comparison with other cellular stresses. Pflügers Archiv - European Journal of Physiology 454, 173–185. URL:http://link.springer.com/10.1007/s00424-006-0195-x, doi:10.1007/s00424-006-0195-x.

[3] Brizi, L., Giampieri, E., Fantazzini, P., Castellani, G., Remondini, D., Zironi, I., 2020. Water exchange between intra and extracellular compartments studied by nuclear magnetic resonance relaxometry and optical microscopy. Journal of Physics D: Applied Physics 53, 085401. URL:https://iopscience.iop.org/article/10.1088/1361-6463/ab538e, doi:10.1088/1361-6463/ab538e.

[4] Burg, M.B., 1995. Molecular basis of osmotic regulation. American Journal of Physiology-Renal Physiology 268, F983–F996. URL:http://www.ncbi.nlm.nih.gov/pubmed/7611465https://www.physiology.org/doi/10.1152/ajprenal.1995.268.6.F983, doi:10.1152/ajprenal.1995.268.6.F983.

[5] Casali, C.I., Weber, K., Favale, N.O., Tome, M.C.F., 2013. Environmental hyperosmolality regulates phospholipid biosynthesis in the renal epithelial cell line MDCK. Journal of Lipid Research 54, 677–691. URL:https://linkinghub.elsevier.com/retrieve/pii/S0022227520419339, doi:10.1194/jlr.M031500.

[6] Delarue, M., Montel, F., Caen, O., Elgeti, J., Siaugue, J.M., Vignjevic, D., Prost, J., Joanny, J.F., Cappello, G., 2013. Mechanical Control of Cell flow in Multicellular Spheroids. Physical Review Letters 110, 138103. URL:https://link.aps.org/doi/10.1103/PhysRevLett.110.138103, doi:10.1103/PhysRevLett.110.138103.

[7] Dolega, M., Zurlo, G., Goff, M.L., Greda, M., Verdier, C., Joanny, J.F., Cappello, G., Recho, P., 2021a. Mechanical behavior of multi-cellular spheroids under osmotic compression. Journal of the Mechanics and Physics of Solids 147, 104205. URL:https://linkinghub.elsevier.com/retrieve/pii/S0022509620304269, doi:10.1016/j.jmps.2020.104205,arXiv:2011.01131.

[8] Dolega, M.E., Delarue, M., Ingremeau, F., Prost, J., Delon, A., Cappello, G., 2017. Cell-like pressure sensors reveal increase of mechanical stress towards the core of multicellular spheroids under compression. Nature Communications 8, 14056. URL:http://www.nature.com/articles/ncomms14056, doi:10.1038/ncomms14056.

[9] Dolega, M.E., Monnier, S., Brunel, B., Joanny, J.F., Recho, P., Cappello, G., 2021b. Extra-cellular matrix in multicellular aggregates acts as a pressure sensor controlling cell proliferation and motility. eLife 10, 1–33. doi:10.7554/eLife.63258.

[10] Ghibaudo, M., Saez, A., Trichet, L., Xayaphoummine, A., Browaeys, J., Silberzan, P., Buguin, A., Ladoux, B., 2008. Traction forces and rigidity sensing regulate cell functions. Soft Matter 4, 1836. URL: http://xlink.rsc.org/?DOI=b804103b, doi:10.1039/b804103b.

[11] Hannezo, E., Prost, J., Joanny, J.F., 2014. Theory of epithelial sheet morphology in three dimensions. Proceedings of the National Academy of Sciences 111, 27–32. URL:http://www.pnas.org/cgi/doi/10.1073/pnas.1312076111, doi:10.1073/pnas.1312076111.

[12] Heinrich, M.A., Alert, R., LaChance, J.M., Zajdel, T.J., Košmrlj, A., Cohen, D.J., 2020. Size-dependent patterns of cell proliferation and migration in freely-expanding epithelia. eLife 9, 1–21. URL: https://elifesciences.org/articles/58945, doi:10.7554/eLife.58945.

[13] Höllring, K., Hubert, M., Smith, A.S., 2023. Substrate Stiffness and Cell Area Predict Cellular Traction Stresses in Single Cells and Cells in Contact. TBD tbd, tbd. URL: tbd, doi: tbd.

[14] Janmey, P.A., Fletcher, D.A., Reinhart-King, C.A., 2020. Stiffness Sensing by Cells. Physiological Reviews 100, 695–724. URL:https://journals.physiology.org/doi/10.1152/physrev.00013.2019, doi:10.1152/physrev.00013.2019.

[15] Kaliman, S., Hubert, M., Wollnik, C., Nuić, L., Vurnek, D., Gehrer, S., Lovrić, J., Dudziak, D., Rehfeldt, F., Smith, A.s., 2021. Mechanical Regulation of Epithelial Tissue Homeostasis. Physical Review X 11, 031029. URL:https://link.aps.org/doi/10.1103/PhysRevX.11.031029, doi:10.1103/PhysRevX.11.031029.

[16] Kaliman, S., Jayachandran, C., Rehfeldt, F., Smith, A.S., 2014. Novel Growth Regime of MDCK II Model Tissues on Soft Substrates. Biophysical Journal 106, L25–L28. URL:https://linkinghub.elsevier.com/retrieve/pii/S0006349514002203, doi:10.1016/j.bpj.2013.12.056.

[17] Lee, S.D., Choi, S.Y., Lim, S.W., Lamitina, S.T., Ho, S.N., Go, W.Y., Kwon, H.M., 2011. TonEBP stimulates multiple cellular pathways for adaptation to hypertonic stress: organic osmolyte-dependent and - independent pathways. American Journal of Physiology-Renal Physiology 300, F707–F715. URL:https://www.physiology.org/doi/10.1152/ajprenal.00227.2010, doi:10.1152/ajprenal.00227.2010.

[18] Lovrić, J., Kaliman, S., Barfuss, W., Schröder-Turk, G.E., Smith, A.S., 2019. Geometric effects in random assemblies of ellipses. Soft Matter 15, 8566–8577. URL:http://xlink.rsc.org/?DOI=C9SM01067J, doi:10.1039/C9SM01067J.

[19] Marel, A.K., Zorn, M., Klingner, C., Wedlich-Söldner, R., Frey, E., Rädler, J.O., 2014. Flow and diffusion in channel-guided cell migration. Biophysical Journal 107, 1054–1064. doi:10.1016/j.bpj.2014.07.017.

[20] McManus, M.L., Churchwell, K.B., Strange, K., 1995. Regulation of Cell Volume in Health and Disease. New England Journal of Medicine 333, 1260–1267. URL:http://www.nejm.org/doi/10.1056/NEJM199511093331906, doi:10.1056/NEJM199511093331906.

[21] Mohammed, D., Park, C.Y., Fredberg, J.J., Weitz, D.A., 2021. Tumorigenic mesenchymal clusters are less sensitive to moderate osmotic stresses due to low amounts of junctional E-cadherin. Scientific Reports 11, 1–12. URL:https://doi.org/10.1038/s41598-021-95740-x, doi:10.1038/s41598-021-95740-x.

[22] Montel, F., Delarue, M., Elgeti, J., Malaquin, L., Basan, M., Risler, T., Cabane, B., Vignjevic, D., Prost, J., Cappello, G., Joanny, J.F., 2011. Stress Clamp Experiments on Multicellular Tumor Spheroids. Physical Review Letters 107, 188102. URL:https://link.aps.org/doi/10.1103/PhysRevLett.107.188102, doi:10.1103/PhysRevLett.107.188102.

[23] Murrell, M.P., Voituriez, R., Joanny, J.F., Nassoy, P., Sykes, C., Gardel, M.L., 2014. Liposome adhesion generates traction stress. Nature Physics 10, 163–169. URL:http://www.nature.com/articles/nphys2855, doi:10.1038/nphys2855.

[24] Narayanan, V., Schappell, L.E., Mayer, C.R., Duke, A.A., Armiger, T.J., Arsenovic, P.T., Mohan, A., Dahl, K.N., Gleghorn, J.P., Conway, D.E., 2020. Osmotic Gradients in Epithelial Acini Increase Mechanical Tension across E-cadherin, Drive Morphogenesis, and Maintain Homeostasis. Current Biology 30, 624–633.e4. URL:https://doi.org/10.1016/j.cub.2019.12.025, doi:10.1016/j.cub.2019.12.025.

[25] Nikpey, E., Karlsen, T.V., Rakova, N., Titze, J.M., Tenstad, O., Wiig, H., 2017. High-salt diet causes osmotic gradients and hyperosmolality in skin without affecting interstitial fluid and lymph. Hypertension 69, 660–668.

[26] Pajtók, C., Veres-Székely, A., Agócs, R., Szebeni, B., Dobosy, P., Németh, I., Veréb, Z., Kemény, L., Szabó, A.J., Vannay, Á., et al., 2021. High salt diet impairs dermal tissue remodeling in a mouse model of imq induced dermatitis. PloS one 16, e0258502.

[27] Paluch, E., Heisenberg, C.P., 2009. Biology and Physics of Cell Shape Changes in Development. Current Biology 19, R790–R799. URL:https://linkinghub.elsevier.com/retrieve/pii/S0960982209014511, doi:10.1016/j.cub.2009.07.029.

[28] Pelham, R.J., Wang, Y.l., 1997. Cell locomotion and focal adhesions are regulated by substrate flexibility. Proceedings of the National Academy of Sciences 94, 13661–13665. URL:http://www.pnas.org/cgi/doi/10.1073/pnas.94.25.13661, doi:10.1073/pnas.94.25.13661.

[29] Peyton, S.R., Putnam, A.J., 2005. Extracellular matrix rigidity governs smooth muscle cell motility in a biphasic fashion. Journal of Cellular Physiology 204, 198–209. URL:https://onlinelibrary.wiley.com/doi/10.1002/jcp.20274, doi:10.1002/jcp.20274.

[30] Pietuch, A., Brückner, B.R., Janshoff, A., 2013. Membrane tension homeostasis of epithelial cells through surface area regulation in response to osmotic stress. Biochimica et Biophysica Acta (BBA) - Molecular Cell Research 1833, 712–722. URL:http://dx.doi.org/10.1016/j.bbamcr.2012.11.006 https://linkinghub.elsevier.com/retrieve/pii/S0167488912003278, doi:10.1016/j.bbamcr.2012.11.006.

[31] Preibisch, S., Saalfeld, S., Tomancak, P., 2009. Globally optimal stitching of tiled 3D microscopic image acquisitions. Bioinformatics 25, 1463–1465. URL:https://academic.oup.com/bioinformatics/article-lookup/doi/10.1093/bioinformatics/btp184, doi:10.1093/bioinformatics/btp184.

[32] Rana, P.S., Kurokawa, M., Model, M.A., 2020. Evidence for macromolecular crowding as a direct apoptotic stimulus. Journal of Cell Science 133. URL:https://journals.biologists.com/jcs/article/133/9/jcs243931/224841/Evidence-for-macromolecular-crowding-as-a-direct, doi:10.1242/jcs.243931.

[33] Rim, J.S., Atta, M.G., Dahl, S.C., Berry, G.T., Handler, J.S., Kwon, H.M., 1998. Transcription of the Sodium/myo-Inositol Cotransporter Gene Is Regulated by Multiple Tonicity-responsive Enhancers Spread over 50 Kilobase Pairs in the 5’-Flanking Region. Journal of Biological Chemistry 273, 20615–20621. URL:https://linkinghub.elsevier.com/retrieve/pii/S0021925818490989, doi:10.1074/jbc.273.32.20615.

[34] Ritter, M., Paulmichl, M., Lang, F., 1991. Further characterization of volume regulatory decrease in cultured renal epitheloid (MDCK) cells. Pflugers Archiv European Journal of Physiology 418, 35–39. URL:http://link.springer.com/10.1007/BF00370449, doi:10.1007/BF00370449.

[35] Roffay, C., Molinard, G., Kim, K., Urbanska, M., Andrade, V., Barbarasa, V., Nowak, P., Mercier, V., García-Calvo, J., Matile, S., Loewith, R., Echard, A., Guck, J., Lenz, M., Roux, A., 2021. Passive coupling of membrane tension and cell volume during active response of cells to osmosis. Proceedings of the National Academy of Sciences 118, e2103228118. URL:http://www.pnas.org/lookup/doi/10.1073/pnas.2103228118, doi:10.1073/pnas.2103228118.

[36] Sandersius, S.A., Chuai, M., Weijer, C.J., Newman, T.J., 2011. Correlating Cell Behavior with Tissue Topology in Embryonic Epithelia. PLoS ONE 6, e18081. URL:https://dx.plos.org/10.1371/journal.pone.0018081, doi:10.1371/journal.pone.0018081.

[37] Schramek, H., Gstraunthaler, G., Willinger, C.C., Pfaller, W., 1993. Hyperosmolality regulates endothelin release by Madin-Darby canine kidney cells. Journal of the American Society of Nephrology 4, 206–213. URL:https://jasn.asnjournals.org/lookup/doi/10.1681/ASN.V42206, doi:10.1681/ASN.V42206.

[38] Sheikh-Hamad, D., Gustin, M.C., 2004. MAP kinases and the adaptive response to hypertonicity: functional preservation from yeast to mammals. American Journal of Physiology-Renal Physiology 287, F1102–F1110. URL:https://www.physiology.org/doi/10.1152/ajprenal.00225.2004, doi:10.1152/ajprenal.00225.2004.

[39] Streichan, S.J., Hoerner, C.R., Schneidt, T., Holzer, D., Hufnagel, L., 2014. Spatial constraints control cell proliferation in tissues. Proceedings of the National Academy of Sciences 111, 5586–5591. URL:http://www.pnas.org/cgi/doi/10.1073/pnas.1323016111, doi:10.1073/pnas.1323016111.

[40] Stringer, C., Wang, T., Michaelos, M., Pachitariu, M., 2021. Cellpose: a generalist algorithm for cellular segmentation. Nature methods 18, 100–106. doi:10.1038/s41592-020-01018-x.

[41] Tokuda, S., Hirai, T., Furuse, M., 2016. Effects of Osmolality on Paracellular Transport in MDCK II Cells. PLOS ONE 11, e0166904. URL:https://dx.plos.org/10.1371/journal.pone.0166904, doi:10.1371/journal.pone.0166904.

[42] Tzvetkova-Chevolleau, T., Stéphanou, A., Fuard, D., Ohayon, J., Schiavone, P., Tracqui, P., 2008. The motility of normal and cancer cells in response to the combined influence of the substrate rigidity and anisotropic microstructure. Biomaterials 29, 1541–1551. URL:https://linkinghub.elsevier.com/retrieve/pii/S0142961207010137, doi:10.1016/j.biomaterials.2007.12.016.

[43] Wullkopf, L., West, A.K.V., Leijnse, N., Cox, T.R., Madsen, C.D., Oddershede, L.B., Erler, J.T., 2018. Cancer cells’ ability to mechanically adjust to extracellular matrix stiffness correlates with their invasive potential. Molecular Biology of the Cell 29, 2378–2385. URL:https://www.molbiolcell.org/doi/10.1091/mbc.E18-05-0319, doi:10.1091/mbc.E18-05-0319.

[44] Yamauchi, A., Uchida, S., Kwon, H.M., Preston, A.S., Robey, R.B., Garcia-Perez, A., Burg, M.B., Handler, J.S., 1992. Cloning of a Na(+)- and Cl(−)-dependent betaine transporter that is regulated by hypertonicity. Journal of Biological Chemistry 267, 649–652. URL:https://linkinghub.elsevier.com/retrieve/pii/S0021925818485432, doi:10.1016/S0021-9258(18)48543-2.

[45] Yamauchi, A., Uchida, S., Preston, A.S., Kwon, H.M., Handler, J.S., 1993. Hypertonicity stimulates transcription of gene for Na(+)-myo-inositol cotransporter in MDCK cells. American Journal of Physiology-Renal Physiology 264, F20–F23. URL:https://www.physiology.org/doi/10.1152/ajprenal.1993.264.1.F20, doi:10.1152/ajprenal.1993.264.1.F20.

[46] Yancey, P.H., Clark, M.E., Hand, S.C., Bowlus, R.D., Somero, G.N., 1982. Living with Water Stress: Evolution of Osmolyte Systems. Science 217, 1214–1222. URL:https://www.science.org/doi/10.1126/science.7112124, doi:10.1126/science.7112124.

